# Serum Exosomes from a Uniquely Defined Early-Adult Donor Cohort Reprogram Endothelial Transcriptomes Linked to Alzheimer’s Pathogenesis

**DOI:** 10.1101/2025.11.20.689131

**Authors:** Brian E Packard, Pierre F Roux, Stephen C L’Heureux, Zamboni William C. Zamboni

## Abstract

Age-related changes in circulating exosomes are implicated in cerebrovascular aging and the pathogenesis of Alzheimer’s disease (AD). Neurovascular dysfunction and blood-brain barrier (BBB) breakdown are recognized as early events in AD, often preceding amyloid-β deposition. Primary human brain micro endothelial cells (HBMECs) from a 38-year-old male were treated with exosomes from young (18–25 years) and old (65–72 years) donors. Whole transcriptomic RNA sequencing analysis identified 5,432 differentially expressed genes, which were organized into five transcriptional clusters. Two principal clusters demonstrated reciprocal patterns: 1) exosomes derived from serum of older adult donors (65–72 years) downregulated genes essential for mitochondrial function (e.g., oxidative phosphorylation) and protein synthesis (e.g., ribosomal biogenesis) and 2) upregulating genes linked to inflammation, junctional remodeling, and proliferative signaling. Crucially, subsequent treatment with exosomes derived from serum of young adult donors (18–25 years) reversed these detrimental transcriptomic profiles via restoration of the expression of mitochondrial and ribosomal machinery toward baseline and suppressed the inflammatory and maladaptive proliferative signaling induced by exosomes derived from serum of older adult donors. These findings demonstrate that exosomes derived from serum of young adult donors can counteract detrimental signals of aging at the transcriptional level, reinforcing the cellular architecture underlying BBB integrity. This supports the therapeutic potential of using exosomes derived from serum of young adult donors to reverse endothelial aging and interrupt the early neurovascular dysfunction that contributes to the progression of AD.

## Introduction

Neurovascular dysfunction and blood–brain barrier (BBB) breakdown are increasingly recognized as initiating events in the pathogenesis of Alzheimer’s disease (AD). Cerebrovascular alterations, including endothelial impairment, reduced cerebral blood flow, and BBB leakage, emerge as early abnormalities in AD, preceding the deposition of amyloid-β and the formation of tau pathology ^1,2,3^. Compromise of the BBB permits the influx of plasma proteins, peripheral immune cells, and other circulating factors into the brain parenchyma, thereby amplifying neuroinflammation and accelerating neurodegenerative processes^4,5,6^. Importantly, these vascular abnormalities persist throughout the course of the disease, sustaining a cycle of oxidative stress, mitochondrial dysfunction, and chronic inflammation that further destabilizes the neurovascular unit^7,8,9^. Thus, vascular dysfunction and BBB disruption represent both initiating and perpetuating mechanisms in AD progression, underscoring the central role of the cerebrovasculature in disease development and as a potential target for therapeutic intervention.

In view of the central role of neurovascular dysfunction and BBB breakdown as early and persistent mechanisms in development of AD, interventions that stabilize cerebrovascular integrity hold therapeutic promise. Circulating extracellular vesicles, particularly exosomes, are emerging as key systemic mediators of vascular health. Exosomes carry proteins, lipids, and nucleic acids that regulate endothelial metabolism, oxidative stress responses, and junctional stability^10,11^. Age-related changes in circulating exosomes contribute to impaired BBB function, as exosomes derived from serum of older adult donors (55-64 years) are enriched in pro-inflammatory cytokines, senescence-associated microRNAs, and oxidized lipids that propagate endothelial dysfunction^12,13^. Conversely, exosomes derived from young adult donors deliver regenerative cargo, including antioxidant enzymes and microRNAs such as miR-126 and miR-21, which enhance mitochondrial function, suppress inflammation, and reinforce tight junction assembly^14,15,16^. Experimental studies demonstrate that young systemic factors, in part mediated by exosomal signaling, can restore cerebral blood flow, improve BBB integrity, and promote neurogenesis in aged or AD rodent models^17,18,19^. These observations position exosomes as both biomarkers of systemic aging and potential therapeutic vectors capable of counteracting vascular decline and reversing BBB dysfunction associated with AD. Exosomes are nanoscale extracellular vesicles of 30–150 nm in diameter that circulate in the bloodstream and mediate intercellular and extracellular communication by transmitting proteins, lipids, and nucleic acids^20^ (Fig.1). Exosomes derived from young organisms have demonstrated rejuvenating effects in aged tissues by delivering proregenerative cargo, including specific microRNAs and antioxidant enzymes^21.22,23,24,25.26,27^. Conversely, exosomes from older donors often carry pro-inflammatory factors, senescence-associated microRNAs, and oxidized lipids that propagate aging phenotypes in recipient cells^28,29,30,31,32^. These observations suggest that exosome age may play a pivotal role in modulating endothelial phenotype and BBB function across the lifespan.

**Fig. 1.**
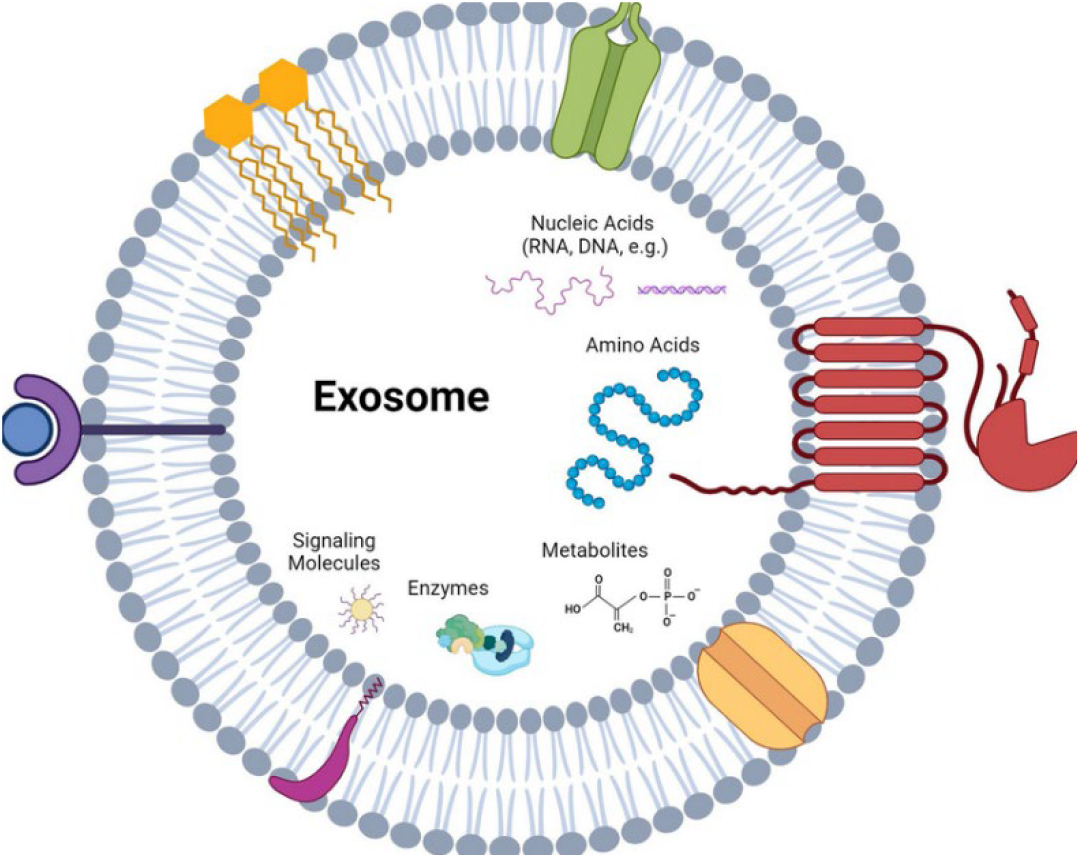
The principal molecular constituents of an exosome, including nucleic acids, proteins, lipids, metabolites, amino acids, signaling molecules, and enzymes. These cargo classes collectively represent the diverse bioactive components packaged within exosomes and underscore their capacity to mediate intercellular communication and functional modulation in recipient cells.

This preclinical study was designed to characterize how exosomes in serum, isolated from young adult and older adult human donors, alter the transcriptome of primary human brain micro endothelial cells (HBMECs). HBMECs are the specialized endothelial cells that line cerebral microvessels and form the principal cellular component of the blood–brain barrier (BBB). They regulate molecular and cellular trafficking between the bloodstream and the brain, maintain cerebral homeostasis, and serve as key mediators of neurovascular function.^33^By identifying differentially expressed genes (DEGs) and the pathways they represent, this study elucidates the mechanisms through which exosomal donor age influences endothelial cell function and to assess whether young (18–25 years) exosomes could reverse the deleterious effects of exosomes derived from serum of older adult donors (65–72 years). The findings have implications for exosome-based therapeutic approaches targeting early neurovascular impairments associated with aging and the development of AD.

## Materials and Methods

### Exosome Isolation

Serum samples (n=3 per cohort) were obtained from healthy human donors grouped by age: young donors (18– 25 years) and old donors (65–72 years) (BioIVT, Product# HUMNSRM-0101025). For exosome isolation, 20 ml of serum was processed to isolate the exosomes using affinity purification kit (Plasma/Serum Exosome Purification Kit, Part# 57600) according to the kit manufactures protocol. The concentration and size distribution of isolated vesicles was assessed via nanoparticle tracking analysis (NanoSight instrument, UNC Nanoparticle Facility, Chapel Hill, NC).

### Cell Culture Studies

For all studies, Human Brain micro-Endothelial cells (HBMEC’s), isolated from a 38-year-old male donor, (Neuromics Inc., Product Code: HEC02) were used. Upon receipt to our facility, cells were thawed and seeded at a density of 1 × 10^5^ cells per well in 48 well tissue culture plates and cultured to confluence in plating media (DMEM/F12 with 10% FBS, endothelial growth supplements) for 24 hours. Four experimental conditions were established in the treatment of the cells, each in biological triplicate: (1) a control group of untreated HBMECs (DMEM/F12 with 3% exosome-depleted human serum); (2) a young exosomes derived from young donors (18–25 years) at a concentration of 1 × 10^12^ exosomes per ml for 24 hours; (3) an old exosomes derived from older donors (65– 72 years) at 1 × 10^12^ exosomes per ml for 24 hours; and (4) an “old-then-young” group in which cells first received old exosomes at 1 × 10^12^ exosomes per ml for 24 hours and subsequently replaced with young exosomes at 1 × 10^12^ exosomes per ml for an additional 24 hours. Media for all conditions was replaced every 8 hours. Following incubation, media was removed and cells were washed with 1 ml of PBS two times. Plates were sealed and frozen at -80°C.

### Total RNA Isolation

Total RNA, including small RNAs, was extracted from HBECs using TRIzol reagent, and RNA integrity was confirmed by Bioanalyzer, with all samples exhibiting RIN values of 8 or higher. Polyadenylated RNA libraries were prepared using the TruSeq RNA Library Prep Kit and sequenced on an Illumina NovaSeq 6000 platform to achieve at least 20 million paired-end reads (2 × 100 bp) per sample (Azenta Biotech). Data Analysis Raw sequencing reads underwent quality control analysis with FastQC, and adapter sequences were trimmed using Trim Galore. Reads were aligned to the human reference genome GRCh38 with the STAR aligner. Gene-level quantification was performed using the annotation. Differential expression analysis was conducted using the edgeR package: counts were normalized by the trimmed mean of M-values method, and differentially expressed genes were defined by a false discovery rate below 0.05 and an absolute log_2_ fold change exceeding one. Hierarchical clustering based on one minus Pearson correlation with Ward.D2 linkage was used to partition DEGs into clusters. Pathway enrichment analysis for KEGG and Hallmark gene sets employed hypergeometric testing, with significance thresholds set at p < 0.01; enrichment results are reported as −log_10_ (p-value).

## Results

### Global Transcriptomic Shifts Induced by Donor-Age Exosomes

A total of 5,432 genes met differential expression criteria (FDR < 0.05) across the control, young, old and sequential treatment of old then young groups. Hierarchical clustering of these genes yielded five distinct clusters, revealing donor-age–dependent transcriptional programs (Figure 2A). Among these, Clusters 4 and 5 displayed the most pronounced reciprocal patterns. Cluster 5 genes were strongly suppressed by old exosomes and restored by young exosomes, whereas Cluster 4 genes were upregulated by old exosomes and reduced toward baseline by young exosomes (Figure 2B, 2C). These large-scale shifts suggest that exosomes carry age-defined cargo capable of profoundly reprogramming endothelial transcriptomes.

**Figure 2.**
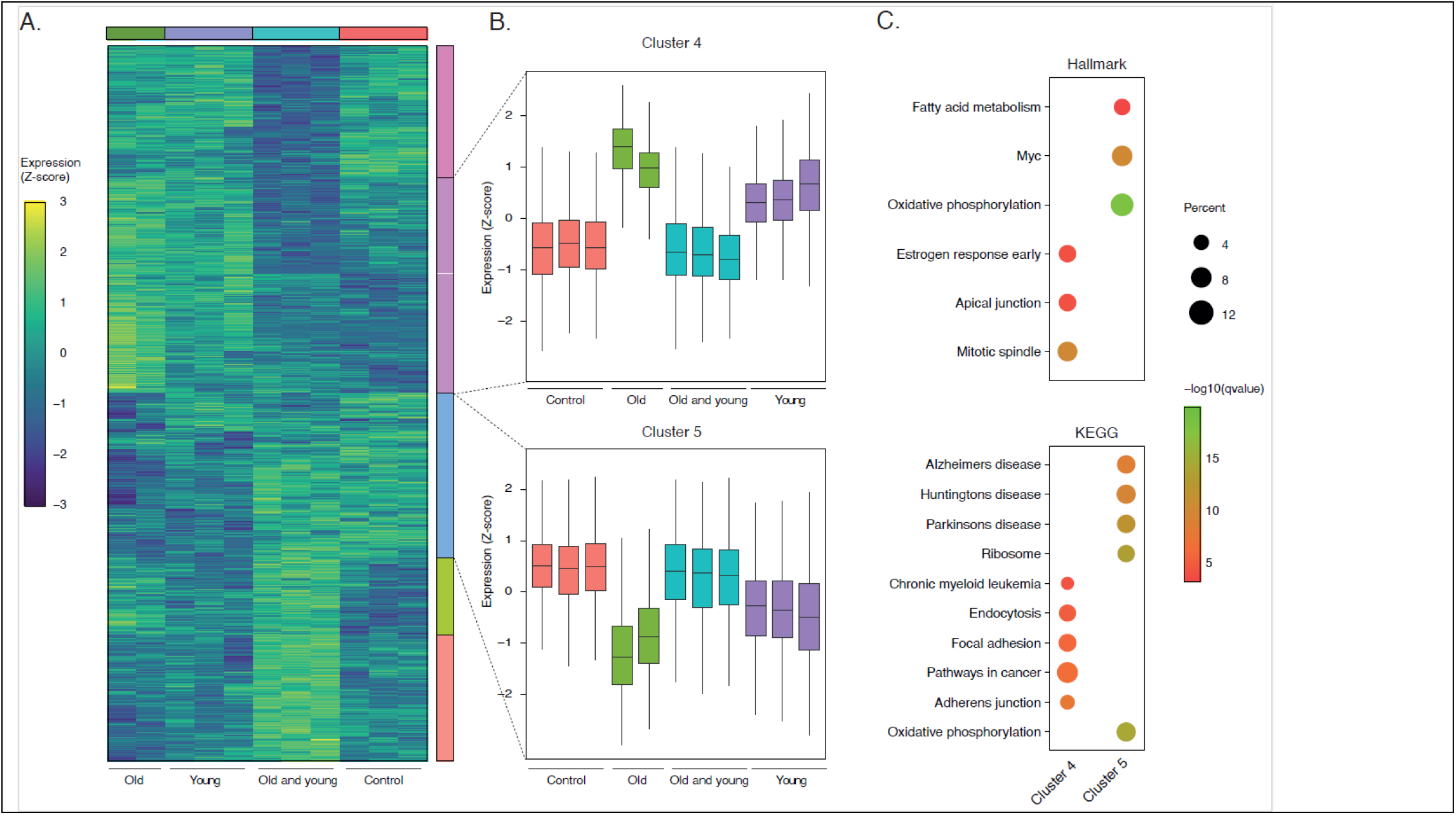
Global transcriptomic reprogramming of HBMECs by serum exosomes from early-adult versus older donors. 2A. A heatmap of results in cells was generated using the top DEGs (n = 5432, filtered for significance) to visualize transcriptional shifts. Expression values were scaled by gene (row-wise Z-scores). Genes were grouped into five clusters using hierarchical clustering based on Pearson’s correlation distance and Ward.D2 linkage, applied to the gene expression matrix. 2B. The heatmap highlights two prominent transcriptional programs associated with exosome exposure: Cluster 4 includes genes <u>upregulated</u> upon exposure of cells to old exosomes, with expression profiles that revert toward baseline (i.e., Control) levels in cells treated with young exosomes alone and old-then-young exosomes. Cluster 5 includes genes <u>downregulated</u> in cells treated with old exosomes that revert toward baseline (i.e., Control) levels in cells treated with young exosomes alone and old-then-young exosomes. 2C. KEGG and Hallmark pathway over-representation of the two most biologically relevant clusters. Color intensity reflects the statistical significance of enrichment, plotted as −log_10_(p-value); darker colors indicate more significant enrichment. Circle size indicates the proportion of overlap between genes in the cluster and genes in the pathway; larger circles represent greater overlap

### Exosomes from Older Donors Suppress Mitochondrial and Translational Programs

Cluster 5 analysis revealed that old exosomes significantly downregulated genes required for mitochondrial oxidative phosphorylation, fatty acid metabolism, and ribosomal biogenesis (Figure 2B). Specific targets included core components of the electron transport chain (NDUFS3, COX4I1, ATP5F1A), metabolic enzymes (CPT1A, IDH3B, PDHB), and cytosolic as well as mitochondrial ribosomal proteins. These changes reflect impaired mitochondrial respiration, reduced ATP production, and diminished protein synthesis capacity— molecular hallmarks of endothelial and vascular aging. Notably, these alterations overlap with pathways previously linked to cerebrovascular decline in Alzheimer’s disease.

### Old Exosomes Drive Inflammatory and Junctional Remodeling

Cluster 4 encompassed genes upregulated by aged exosome treatment, consistent with an endothelial stress response (Figure 2B). Junctional proteins (CDH1, CTNNA1, COL4A1, laminins), inflammatory mediators (MMP9, ICAM1, VEGFB), and proliferative signaling molecules (EGFR, ERBB2, PIK3CB, MTOR) were strongly induced. These changes suggest maladaptive remodeling of cell–cell junctions and activation of oncogenic and inflammatory pathways. Such remodeling compromises barrier integrity and promotes a pro-inflammatory endothelial phenotype—features consistent with the “inflammaging” signature of aged vasculature. Importantly, these transcriptomic changes highlight how aged exosomal cargo may directly destabilize blood–brain barrier function.

### Young Exosomes Restore Mitochondrial and Ribosomal Gene Expression

In contrast, treatment with young exosomes either maintained expression of mitochondrial and translational machinery close to control levels or reversed the suppressive effects of old exosomes. Mitochondrial respiratory chain subunits, ribosomal proteins, and metabolic transporters exhibited partial or complete restoration toward baseline (Figure 2C). This indicates that young exosomal cargo actively supports energy metabolism, protein synthesis, and overall endothelial health, counteracting the decline triggered by aged systemic signals.

### Sequential Treatment with Young Exosomes Reverse Effects of Old Exosomes

When HBMECs were first exposed to old exosomes and subsequently treated with young exosomes, transcriptomic profiles demonstrated broad recovery. Mitochondrial and ribosomal gene expression, initially suppressed by old exosomes, was restored toward baseline, while inflammatory and junctional stress responses were attenuated. The sequential design underscores that young exosomes not only preserve baseline endothelial programs but also exert restorative effects after old-induced dysfunction has already been established. This highlights their potential therapeutic application in reversing age-associated endothelial injury.

### Pathway Enrichment Highlights Neurovascular Mechanisms of Aging and Reversal

Pathway enrichment analyses reinforced these findings, revealing over-representation of oxidative phosphorylation, fatty acid metabolism, ribosomal biogenesis, inflammatory response, and apical junction organization. Old exosomes disrupted mitochondrial and translational pathways while activating inflammatory and proliferative programs, aligning with mechanisms implicated in cerebrovascular aging and Alzheimer’s disease. Conversely, young exosomes restored mitochondrial and translational capacity and suppressed maladaptive junctional and inflammatory signaling. Collectively, these results demonstrate that young exosomes reprogram endothelial transcriptomes to stabilize BBB-associated functions and mitigate neurovascular decline associated with old exosomes.

Pathway enrichment analyses reinforced these findings, revealing over-representation of oxidative phosphorylation, fatty acid metabolism, ribosomal biogenesis, inflammatory response, and apical junction organization. Old exosomes disrupted mitochondrial and translational pathways while simultaneously activating inflammatory and proliferative programs mechanisms strongly implicated in cerebrovascular aging and Alzheimer’s disease pathogenesis. In contrast, young exosomes restored mitochondrial and translational capacity while suppressing maladaptive junctional remodeling and inflammatory signaling. Collectively, these results demonstrate that young exosomes reprogram the endothelial transcriptome to stabilize barrier integrity and mitigate the neurovascular decline induced by aged exosomes.

## Discussion

The results of these studies demonstrate that exosomes from the human serum of old donors (65-72 years) transmit pro-aging signals to primary HBECs, driving transcriptional programs associated with mitochondrial dysfunction, translational decline, inflammatory activation, and maladaptive junctional remodeling. These results are consistent with prior studies showing that exosomes derived from serum of older adult donors are known to carry increased levels of pro-inflammatory cytokines (including interleukin-6 and tumor necrosis factor-α), senescence-associated microRNAs, and oxidized lipids that activate NF-κB and stress-response pathways in recipient cells^34,35^. The upregulation of pro-inflammatory, junctional, and oncogenic genes observed in Cluster 4 closely mirrors the “inflammaging” phenotype described in aged cerebrovascular endothelium, wherein endothelial-to-mesenchymal transition and barrier leakiness contribute to neurovascular compromise^36,37^.

Exosomes from serum of young adult donors (18–25 years), by contrast, exhibited a capacity to reverse these age-induced transcriptomic alterations. Mechanistically, exosomes derived from serum of young adult donors (18–25 years) deliver enriched cargo of microRNAs—particularly miR-126 and miR-21—that target inhibitors of PI3K/Akt and MAPK signaling pathways to enhance mitochondrial biogenesis, upregulate antioxidant defenses through Nrf2 activation, and promote tight junction protein expression. miR- 126 has been shown to facilitate endothelial proliferation and migration under stress conditions^38,39^, thus supporting barrier repair.

Proteomic analyses of exosomes derived from serum of young adult donors (18–25 years) have revealed the presence of antioxidant enzymes (such as superoxide dismutase 1 and catalase) and growth factors (including vascular endothelial growth factor and fibroblast growth factor 2) that further bolster mitochondrial respiration, mitigate reactive oxygen species accumulation, and stimulate pro-survival signaling^40,41^. Differential expression of integrins and other surface ligands on exosomes derived from serum of young adult donors (18–25 years) may also modulate vesicle uptake kinetics and trafficking, preferentially activating regenerative pathways in recipient endothelial cells.

The transcriptomic restoration induced by exosomes derived from serum of young adult donors (18–25 years) holds significant implications for AD. Early neurovascular dysfunction—manifested as endothelial mitochondrial impairment, reduced nitric oxide production, diminished CBF, and BBB breakdown—precedes amyloid deposition and tau pathology^42,43,44,45,46^. Impaired barrier function permits infiltration of plasma proteins and immune cells into the brain parenchyma, fueling neuroinflammation and amyloid accumulation^6,47,48,49,50^. By revitalizing endothelial metabolic and translational capacity, suppressing inflammatory signaling, and reinforcing junctional integrity, exosomes derived from serum of young adult donors (18–25 years) may stabilize the neurovascular unit and interrupt the cascade of events that lead to AD onset and progression. These findings align with emerging AD biomarkers, such as evidence of BBB breakdown in APOE4 carriers, and parallel clinical approaches like plasma exchange therapies (e.g., AMBAR trial) that aim to remove aged factors and introduce rejuvenating ones^51,52^.

## Conclusions

This study demonstrates that serum-derived exosomes from old human donors (65-72 years) elicit transcriptional changes in primary HBMECs consistent with mitochondrial suppression, translational decline, inflammatory activation, and disorganized junctional signaling—hallmarks of cerebrovascular aging and contributors to AD pathogenesis ^[1, 3].^ Crucially, serum-derived exosomes from young donors (18–25 years) reversed these deleterious transcriptomic profiles, restoring gene expression associated with oxidative phosphorylation, ribosomal biogenesis, tight junction assembly, and anti-inflammatory signaling. These findings underscore that age critically influences exosomal cargo, and that this differential cargo in turn determines the function of human brain endothelial cells.

Accordingly, exosome-based therapeutics may offer a non-cellular, scalable strategy to counteract neurovascular decline in aging and AD. To advance toward clinical application, future work should prioritize functional validation, comprehensive exosome characterization and standardization, and rigorous in vivo pharmacology, efficacy, and safety assessments.

## Study Limitations and Future Directions

Notwithstanding these promising results, several limitations warrant consideration. First, the in vitro HBEC model lacks the full cellular complexity of the neurovascular and physiologic structure of the brain, which includes interactions with pericytes, astrocytes, and neurons. Future studies should employ rigorous standardization of exosome isolation, cargo characterization (proteomic and small RNA sequencing), and batch reproducibility is necessary before additional preclinical and clinical translational studies are performed. This characterization should include Western blots for markers like CD63 and CD81 to confirm purity. Second, the effects of exosomes need to be evaluated in more complex in vitro systems, such as co-culture systems or three-dimensional organoid models to better capture cell–cell crosstalk. Third, although transcriptomic data provide strong evidence of molecular restoration, functional validation is essential; measures such as mitochondrial respiration assays (e.g., Seahorse extracellular flux analysis), assessments of barrier integrity (transendothelial electrical resistance and permeability assays), and quantification of inflammatory cytokine secretion must confirm phenotypic reversal. Fourth, in vivo experiments using aged or AD transgenic mouse models are critical to determine the optimal dose, regimen, and route of administration of young exosomes to improve cerebrovascular function, reduce amyloid and tau pathology, and enhance cognitive performance. As part of the safety evaluation in the in vivo studies, the evaluation of the immunogenicity of allogeneic exosomes need to be evaluated.

